# Setting a trajectory for CO_2_ emission reduction in academic research: a case study of a French biophysics laboratory

**DOI:** 10.1101/2023.11.30.569421

**Authors:** Caroline Giuglaris, Jean de Seze

**Affiliations:** UMR 168 Laboratoire PhysicoChimie Curie, Institut Curie, PSL Research University, Sorbonne Université, CNRS, Paris, France

**Keywords:** carbon footprint, academia, decarbonisation

## Abstract

Climate change is a scientifically proven phenomenon caused by anthropic activities, which requires urgent and significant reductions in greenhouse gas emissions. Despite the increasing vocalization of scientists advocating for political action, the issue of the environmental impact of academic research has been neglected for some time. Now, field-dependent initiatives have emerged, such as the non-profit organization My Green Lab, which delivers green certifications to biology and chemistry labs, and institute-dependent programs, such as the Max Planck Sustainability Network. In France, an independent collective was founded in 2019 to address the environmental footprint of academic research following the COP 15 Paris Agreement: Labos 1Point5. Building on their resources and methodology, we have quantified the overall carbon footprint of our biophysics laboratory, considering energy consumption, purchases and travel, for the year 2021. We investigate how this footprint would decrease by 2030 following systemic changes (change in the energy mix, improvements from suppliers), and we propose scenarios based on additional voluntary initiatives to reach a final reduction of -50% compared to the 2021 baseline, following IPCC targets. We have now formed a group of more than 20 colleagues to achieve this goal, emphasizing the importance of collective action. Finally, we provide advice based on our own experience to assist others in addressing the environmental impact of academic research in their respective laboratories.

## 1. Introduction

Fundamental research is and will be impacted by the energy [1, 2], climate and biodiversity crisis we currently face, in at least two different ways. First, modern research is highly dependent on public funding, which starts to set constraints to reduce the environmental impact of public expenses[3]. Second, scientists depend on their suppliers for energy, travel, consumables and instruments, who are themselves facing environmental, energetic and legal constraints. Moreover, environmental awareness is more and more spread among researchers, who want to act and take part in transforming the society we live in. To start reducing the environmental impact of current research, one has first to measure it, and then take decisions and actions to follow a given trajectory of emissions reductions. The standard norm for measuring this impact is the quantification of the carbon footprint, which takes into account all the direct and indirect greenhouse gas emissions due to a specific activity. It reveals both how emission-intensive an activity is, as well as how dependent it is on emission-intensive industries. Many initiatives have been taken around the world to start quantifying the carbon footprint of research laboratories [4, 5] and implement measures to reduce emissions [6–8]. In France, a collective of academics has put together a free online tool to compute the environmental impact of one’s laboratory [9]. It allowed the measurement of the carbon footprint of 834 laboratories (as of July 2023) with a standardised protocol, and has led to several publications about the global picture of the carbon footprint of academia in France [10–12]. Now that the main emission sectors are known, laboratories have to define their strategies for aiming at a strong reduction of the carbon footprint of their research and make concrete decisions to reach them.

Here, we provide a detailed example of such a carbon footprint trajectory, for a biophysics department in Institut Curie, Paris, France. Firstly, we present the detailed carbon footprint of the studied laboratory, which comprises both experimental and theory teams and is composed of 160 people. Second, we propose a strategy to reduce emissions, aiming at a 50% reduction for 2030 compared to 2021, following the consensus of the [13]. To compute this trajectory, we take into account both the reductions that were pledged by our main suppliers, together with voluntary actions that have to be taken within the lab to reach the target. We also discuss how the strategy is organized within this research department, with working groups tackling the most important emission sectors.

## 2. Results

### 2.1. Carbon footprint of Physico-Chimie Curie: a biophysics laboratory with experiments and theory

The specific research laboratory presented here is composed of 14 research teams, with approximately 160 full-time equivalent positions, including permanent and short-term scientists, engineers, technicians, and support staff. To compute the carbon footprint, we used the methodology and emission factors proposed by Labos 1Point5 [12]. We could not directly use the online tool GES 1point5 [14] to compute the emissions due to purchases. Indeed, it required a specific classification of the purchases, standardized by the CNRS (one of France’s national research institutes), to which our laboratory is not affiliated. Thus, specific emission factors for aggregated purchases categories have been estimated for this laboratory, which details can be found in Material and Methods.

**Figure 1 (a)** presents the detailed estimated carbon footprint for the year 2021. It gives a global impact of 605 tCO_2_ for this year, with a carbon footprint of 4 tCO_2_/capita. Interestingly, it is very close to the average carbon footprint of French laboratories [15], and similar to what has been computed for another biophysics laboratory in Paris [16]. As for most research laboratories in France, purchases, including consumables and equipment, represent the first sector of emissions, with approximately two-thirds of the total emissions. Indirect energy consumption (referred to as ’Scope 2’), in our case electricity and heating, comes as the second emission source (20%), and travel as the third (6%). Of note, 2021 has been a year of low international travel compared to 2019, because of the restrictions due to the COVID pandemic in many countries.

**Figure 1.**
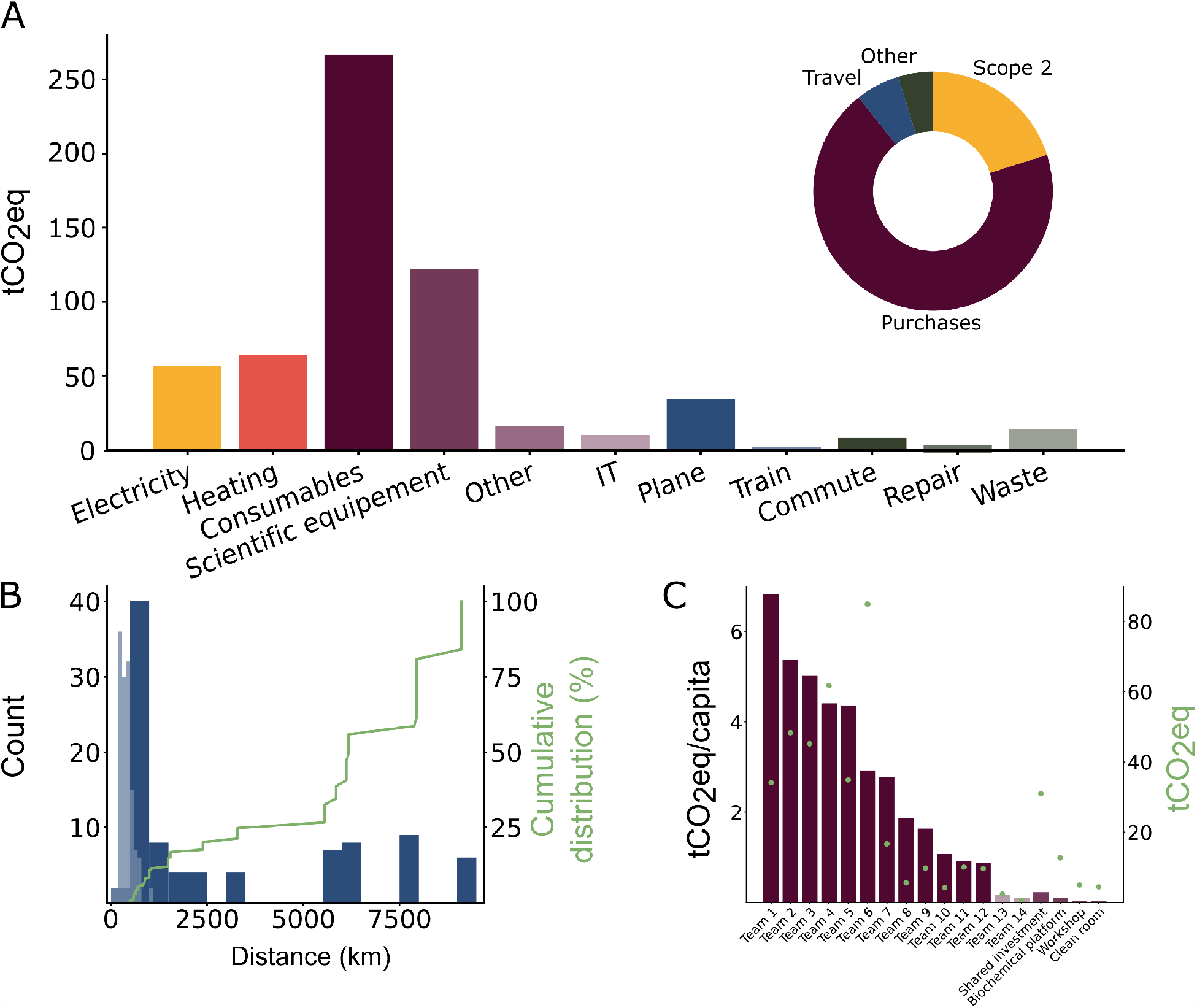
Carbon footprint for the year 2021. a. Carbon footprint of the laboratory with 160 people (in tonnes CO_2_ equivalent) for the main different emission sectors. Top right, pie chart grouping the purchases (in violet), electricity and heating (scope 2, in yellow), travels (in blue), and the other sectors (in green). b. Number of travels depending on the distance travelled for the train (light blue) and the plane (dark blue). In green, cumulative distribution of total kilometres travelled. c. Emission per capita due to purchases for each team of the Department, in decreasing order. In green, total value of the emissions of purchases for each team, right axis.

Purchases are the main source of emissions but also the most specific to the type of research. Our research unit is composed of both experimental and theory teams, with highly different research topics, there is a huge heterogeneity in the emission per capita in each team, as can be seen in **Figure 1 (c)**. It ranges from 7tCO_2_/capita/year for experimental teams, to less than 1 tCO_2_/capita/year for teams doing only theory.

For travels, we detailed in **Figure 1 (b)** the numbers of travels depending on the distance, and the cumulative distance. As already known and discussed multiple times [17], plane, taken mostly for conferences, is by far the main contributor to the impact of travel. As plane emissions are considered to be proportional to the distance travelled, few long-distance travels contribute to most of the CO_2_ emissions for travel.

Overall, the detailed carbon footprint presented here reveals the main sources of emission, as well as the heterogeneity of each of these sources, with some actions and types of research being responsible for a large majority of the emissions. It allows for prioritisation of the actions and construction of a short and long-term strategy for carbon footprint reduction.

### 2.2. Building a trajectory to reduce CO2 emissions

The carbon footprint of the laboratory revealed how much our research is dependent on external suppliers, such as the companies from which we buy our material, our electricity supplier, or plane companies for travelling. As a consequence, the evolution of the carbon footprint of our research unit will be highly dependent on the evolution of the carbon footprint of our suppliers in the coming years. These suppliers have themselves taken pledges for emission reductions, that we need to take into account into our trajectory. Few companies have a clear and realistic trajectory beyond 2030. Thus, we have restricted ourselves to the 2021-2030 period, for both systemic and voluntary changes. As a target, we chose to aim at a 50% reduction for 2030 compared to 2021.

To estimate the emission reduction that would happen without any changes from our side - neither increasing nor decreasing our budget, purchases, travel, electricity or heating consumption - we have based our work on the commitments made by our major suppliers. Details can be found in the Methods sections, and have led us to an estimate of a 23% reduction due to systemic changes. Most of it comes from the reduction in emissions due to purchases (- 30%), electricity (- 17.5%) and planes (- 23%). We did not evaluate the feasibility of the commitments but took the pledges as granted.

To reach a 50% reduction in 2030 compared to 2021, there is still a part that must be achieved through voluntary action - approximately 34%, which is equivalent to 5% each year. We have built a working scenario, with the reduction details for the major sources of carbon emission, presented in **Figure 2 (b)**. This scenario takes into account a mix of decisions at the level of Institut Curie (about 2000 scientists) and improvements in lab-level management. At the level of the institute, renovation of the building should be the measure with the most impact, but also the most costly (- 60% for a cost of at least 1 to 2 Me). At the level of the lab, we opted for a 30% reduction in purchases, a 50% reduction in travel, and a 30% reduction in our electricity consumption.

**Figure 2.**
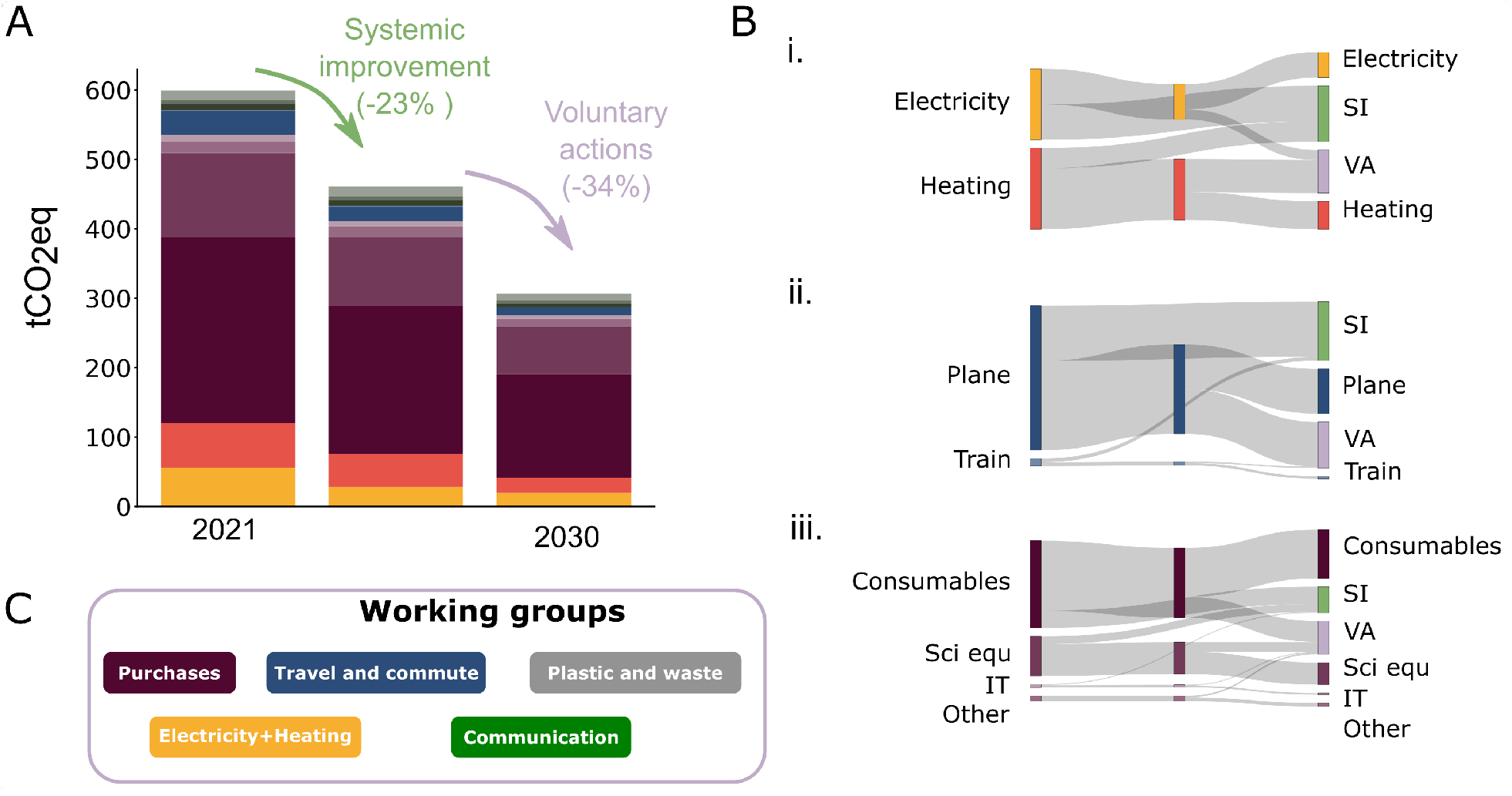
Proposed trajectory. a. Proposed trajectory of emission reduction. Taking 2021 as a reference (left bar), systemic improvements due to suppliers will account for around 23% (middle bar), and voluntary actions need to decrease emissions by 34% to reach a total of 50% reduction (right bar). Colours represent the different sectors presented in 1 b. Detailed reductions, due to systemic improvements (SI) or voluntary actions (VA) for electricity and heating (i), purchases (ii) and travels (iii) c. Working groups created in our department to tackle the different emission sectors.

In order to convert these goals into concrete actions and pathways, we organised into working groups, which focused on the main sub-targets that can be tackled at the level of the laboratory (**Figure 2 (c)**). They refine the computation of the carbon footprint, evaluate the impact of the different decisions that can be taken, and do a follow-up on their emission source. We added a communication team, absolutely required to raise awareness among researchers of the laboratory.

Some actions will be directly measurable, such as reductions in electricity or heating consumption. Other, such as changes in consumption habits, will be hardly visible at a constant budget, because of the use of factors of conversion between euros and CO_2_. Thus, we decided to quantify these actions independently and subtract them every year for a quantified estimate of our actions.

## 3. Discussion

We wanted to present here, as an example, the carbon footprint of a specific biophysics laboratory, mixing experimentalists and theoreticians, and a proposal to build a trajectory for the reduction of the carbon footprint. We showed here that some part should be achieved by the systemic societal changes, but will not be enough to reach the ambitious target of a 50% reduction proposed by the European Union. Reaching such a target will require both institutional and lab-level changes, which may have impacts on research efficiency and quality. These impacts are hard to evaluate, as they could come together with changes in terms of the price of energy or purchases.

One point that we also wish to raise, is the goal that has to be taken for research. Should research do more, or less, than the rest of society?[14] We decided to set, as an arbitrary target, the goal of a 50% reduction for 2030 compared to 2021, which follows the global aim at the scale of the European Union. One could argue that research should be exempted from these efforts. However, we personally think that research should do its part and will anyway have to adapt to the global changes, the sooner, the better. In any case, the strategy used here can be adapted to any target, taking into account systemic and voluntary changes for emission reduction trajectory.

As for all carbon footprints, most values taken here are subject to discussion. We chose the values given by Labos 1Point5, as a point of reference, because they are widely used in the French community. Compared to other laboratories in other countries, the emission factors will be most diverging for the impact of electricity, which is dependent on the electricity mix of the country. France’s electricity mix is low-carbon compared to most countries and gives us a particularly low impact on electricity consumption. In other laboratories, electricity can be the first source of emission. Another impact negligible here compared to other laboratories is the commuting section. Indeed, the laboratory is situated at the heart of Paris, few people need to use their car on a daily basis, which considerably reduces the impact of commute.

We have good reasonable orders of magnitude but are surely not exhaustive on our carbon footprint. One of the unknowns is the impact of external servers used for computation. We did not add any estimate, as most researchers were doing their computations locally - which is then taken into account in the electricity consumption - and we had no way to evaluate the amount and impact of this computation. We don’t have any estimate for refrigerant gases either, which can be strong greenhouse gases but are harder to quantify.

Additionally, this study is focused on the carbon footprint of our laboratory, but we did not assess yet other environmental impacts of our research. Water usage (for material sterilization) and environmental pollution (due to the incineration of bio-contaminated products) are expected to be high and to have a negative impact on biodiversity, air quality etc.

Overall, our study proposes an example of carbon footprint computation and building of a trajectory for emission reduction, which underlines the importance of quantification for targeting the most impactful emission sectors. We think that starting such a project in a laboratory also helps raise awareness among researchers, who can then be involved in their personal lives, or even change their research subject to help our society facing the environmental crisis we are living in.

## 4. Methods

### 4.1. Data collection

To compute the carbon footprint, we used the same methodology proposed by Labo1Point5 in their open-source tool [14]. We used their tool to compute the carbon footprint due to commute, after a survey among the laboratory. We didn’t use the tool for the other categories, as our purchases were not classified through the CNRS inventory numbers. We used macro emission factors - presented in the next subsection - for the emission due to the purchases, and the following ones for the other categories in **Table 1**.

**Table 1:** Emission factors for electricity, heating (district heating) and transportation.

| Category | Emission factor | Source |
| --- | --- | --- |
| Electricity | 55 gCO<sub>2</sub>/kWh | https://www.rte-france.com/eco2mix |
| Heating | 161 gCO<sub>2</sub>/kWh | https://www.cpcu.fr/wp-content/uploads/2020/02/CHIFFRES-CLES-CPCU-2019-sans-plan.pdf |
| Plane | 120 gCO<sub>2</sub>/km | https://www.transitionpathwayinitiative.org/publications/101.pdf?type=Publication |
| Train | 33gCO<sub>2</sub>/km | https://www.eea.europa.eu/publications/rail-and-waterborne-transport |
| Bus | 80 gCO<sub>2</sub>/km | https://www.eea.europa.eu/publications/rail-and-waterborne-transport |

**Table 2:** Emission factors of purchases.

| Category | Macro emission factor |
| --- | --- |
| Lab consumables | 0.74 gCO<sub>2</sub>/€ |
| Scientific equipment | 0.26 gCO<sub>2</sub>/€ |
| Small scientific equipment | 0.30 gCO<sub>2</sub>/€ |
| Administrative and IT supplies | 0.22 gCO<sub>2</sub>/€ |
| Repair and maintenance | 0.14 gCO<sub>2</sub>/€ |
| Other equipment | 0.26 gCO<sub>2</sub>/€ |
| Non-scientific general supplies | 0.27 gCO<sub>2</sub>/€ |
| Other general spendings | 0.26 gCO<sub>2</sub>/€ |
| Other lab spendings | 0.31 gCO<sub>2</sub>/€ |

### 4.2. Macro emission factors for purchases

For the emissions due to purchases, we used the categories existing in our laboratory for orders: Consumables, Big scientific equipment, Small scientific equipment, IT, Repair, Other. Labelling manually 300 hundred items in the CNRS categories - which have known emission factors-, we extrapolated macro emission factors for each category. It gave us the following emission factors, in **Table 4.2**.

### 4.3. Evaluation of systemic changes

To build a scenario of CO_2_ emissions reduction, we first estimated the decrease due to our suppliers’ efforts. To do this, we looked at reports from our main suppliers and assumed that they would follow their own pledges of emissions reduction. From this, we could extrapolate the systemic reduction per category, and design our own working scenario. The numbers and the links to the pledges taken by the main suppliers are in appendix A.

## Acknowledgements

We thank Labo1Point5 for all the work done that helped us in doing this study, and especially Andre Estevez-Torres for the help with the emission factors for purchases. We thank the whole Green Physics Team of UMR168 for its motivation in making things move. We would like to thank all the administration team who helped us in gathering the data, especially Fabrice Demarthon. We would like also to thank Pascal Hersen for his support.

## Appendix A. Scenario

### Appendix A.1. Suppliers’ claims (table)

Appendix A.1. Suppliers' claims (table)
| Supplier | Decrease in 2030 | Source |
| --- | --- | --- |
| Thermofisher | 30% | https://static.fishersci.eu/content/dam/fishersci/en_EU/environment/14654_Sustainability/2019_Corporate_Social_Responsiblity_Report_V2.pdf |
| Merk | 30% | https://www.merckgroup.com/content/dam/web/corporate/images/country-specifics/france/entreprise/rse/strategie_rse_website/home/RSE\%202020\%20PDF.pdf |
| Avantor | 15% | https://www.avantorsciences.com/us/en/about/innovation-environment |
| Nikon | 16% | https://www.nikon.com/company/sustainability/report/2019/sr2019_all.pdf |
| Lenovo | 25% | https://investor.lenovo.com/en/sustainability/reports/FY2022-lenovo-sustainability-report.pdf |
| Electricity | 17.5% | https://assets.rte-france.com/prod/public/2021-04/Bilan\%20previsionnel\%202021.pdf |
| Plane | 23% | https://iea.blob.core.windows.net/assets/deebef5d-0c34-4539-9d0c-10b13d840027/NetZeroBy2050-ARoadmapfortheGlobalEnergySector_CORR.pdf |

## Notes

### Competing Interest Statement

The authors have declared no competing interest.

